# Circulating low-molecular-weight (poly)phenol metabolites in the brain: unveiling in vitro and in vivo blood–brain barrier transport

**DOI:** 10.1101/2024.02.27.582339

**Authors:** Rafael Carecho, Daniela Marques, Diogo Carregosa, Domenico Masuero, Mar Garcia-Aloy, Federica Tramer, Sabina Passamonti, Urska Vrhovsek, M. Rita Ventura, Maria Alexandra Brito, Cláudia Nunes dos Santos, Inês Figueira

## Abstract

Circulating metabolites resulting from colonic metabolism of dietary (poly)phenols are highly abundant in the bloodstream, though still marginally explored, particularly concerning their brain accessibility. Our goal is to disclose (poly)phenol metabolites’ blood–brain barrier (BBB) transport, in vivo and in vitro, as well as their role at BBB level. For three selected metabolites, benzene-1,2-diol-3-sulfate/benzene-1,3-diol-2-sulfate (pyrogallol-sulfate – Pyr-sulf), benzene-1,3-diol-6-sulfate (phloroglucinol-sulfate – Phlosulf), and phenol-3-sulfate (resorcinol-sulfate – Res-sulf), BBB transport was assessed in human brain microvascular endothelial cells (HBMEC). Their potential in modulating in vitro BBB properties at circulating concentrations was also studied. Metabolites’ fate towards the brain, liver, kidney, urine, and blood was disclosed in Wistar rats upon injection. Transport kinetics in HBMEC highlighted different BBB permeability rates, where Pyr-sulf emerged as the most in vitro BBB permeable metabolite. Pyr-sulf was also the most potent regarding BBB properties improvement, namely increased beta(β)-catenin membrane expression and reduction of zonula occludens-1 membrane gaps. Whereas no differences were observed for transferrin, increased expression of caveolin-1 upon Pyr-sulf and Res-sulf treatments was found. Pyrsulf was also capable of modulating gene and protein expression of some solute carrier transporters. Notably, each of the injected metabolites exhibited a unique tissue distribution in vivo, with the remarkable ability to almost immediately reach the brain.

## Introduction

The incidence of chronic neurodegenerative disorders is increasing, with limited therapeutic options that only alleviate patients’ symptoms.^1,2^ Drug-limited brain access, allied to multifactorial pathobiology, renders the discovery of new therapies/preventive approaches critical to efficiently tackle such disorders. (Poly)phenols, abundant in fruits and vegetables, have been extensively studied and emerged as relevant pleiotropic compounds, having the capacity to tackle multiple cellular and molecular mechanisms,^3^ beneficial towards complex multifactorial pathologies, as neurodegenerative ones.^4,5^ Therefore, (poly)phenols comprise very promising potential nutritional/ nutraceutical entities in the prevention of such increasingly prevalent and uncurable disorders. Indeed, greater intake of (poly)phenol-rich foods has been associated with slower rates of cognitive decline, improved memory and learning performance, and enhanced neuronal responses in humans.^6–8^ Recognizing which (poly)phenols or corresponding metabolites found in circulation are responsible for the observed outcomes, and by which mechanisms, however, is still marginally understood. Notably, after ingestion, only 5–10% of the parent/dietary (poly)phenols present in food are absorbed in the small intestine, whereas the great majority reach the colon being extensively metabolized by the gut microbiota into low-molecular weight (LMW) (poly)phenol metabolites.^9,10^ Previous work from our team showed that some LMW (poly)phenol metabolites, namely simple phenolic sulfates, are abundant in circulation after the consumption of a (poly)phenol-rich meal, reaching concentrations between nanomolar to low micromolar in healthy subjects.^11^ Importantly, it has already been shown that phenolic sulfates can arise from different (poly)phenol rich sources, regardless of crops, regions and, season,^10^ which makes them ideal candidates to be explored for their pleiotropic health benefits. But to exert their role in the brain, LMW metabolites need to either transpose the blood–brain barrier (BBB) or be effective at the borders of the BBB, in relevant concentrations comparable to those found in the bloodstream or to the ones reported to be transported across the BBB.^12^ Although brain accessibility of LMW (poly)phenol metabolites might not be compulsory for their benefits,^13^ the mechanisms by which LMW (poly)phenol metabolites can overcome the BBB are still quite underexplored. Passive permeation and carrier-mediated transport (CMT) constitute potential routes by which these molecules could cross the BBB. Twenty years ago, Youdim and co-workers reported the mechanisms of permeation of some dietary (poly)phenols through the BBB, not their circulating metabolites, being linked to their lipophilicity.^14,15^ Nevertheless, other mechanisms should also be considered, like paracellular and vesicular transport.^16^ Studies with parent (poly)phenols suggest that they can be transported across endothelium and epithelium with the involvement of ATP-binding cassette (ABC) transporters, organic anion transporters (OAT), and also organic-anion-transporting polypeptides (OATP) families.^17,18^ For some LMW (poly)phenol metabolites as phenolic acids, interactions with OAT have already been demonstrated.^19,20^ Nevertheless, it is not yet completely clear whether the primary route by which LMW (poly) phenol metabolites reach the brain is by simple diffusion, transcellular transport, or via specific transporters. We have provided evidence that some bioavailable phenolic sulfates are indeed transported across the BBB^12^ and were detected in the cerebrospinal fluid (CSF) of healthy subjects.^21^ Importantly, despite presenting different BBB permeabilities, such LMW (poly)phenol metabolites do not appear to be substrates of main ABC transporters.^12^ Moreover, in silico analysis suggested that none of these would be able to cross the BBB endothelium solely by passive permeation, thus active transport might be involved.^12^ Remarkably, pre-treatment with these BBB permeable phenolic sulfates, in circulating concentrations, improved cellular responses to excitotoxicity and oxidative and inflammatory injuries, particularly via modulation of NF-κB pathway^12^ and two of them emerged as strong attenuators of neurodegeneration against a dopaminergic insult in a three dimensional neuronal model.^22^ Such observations support our hypothesis that LMW (poly)phenol metabolites can be the true responsible for the reported brain health benefits arising from (poly)phenol-rich diets. The exact mechanisms by which they cross BBB and reach the brain, as well as their role at BBB endothelial level, however, is yet to be clarified. Our goal is to disclose circulating LMW (poly)phenol metabolites’ capability of crossing the BBB in vitro and reaching the brain in vivo. To this end, three bioavailable phenolic sulfates, benzene-1,2-diol-3-sulfate/benzene-1,3-diol-2-sulfate (pyrogallol-sulfate isomers, Pyr-sulf), benzene-1,3-diol-6-sulfate (phloroglucinol-sulfate, Phlo-sulf) and phenol-3-sulfate (resorcinol-sulfate, Res-sulf) were chosen to be tested in physiologically relevant concentrations both in an established cellular model of the BBB^12,23^ and injected in Wistar rats mimicking a post-absorption scenario.^24^ Pyr-sulf demonstrated superior in vitro BBB permeability compared to other metabolites. It increased β-catenin expression, reduced zonula occludens (ZO)-1 gaps, and upregulated caveolin-1 (Cav1), supporting its potential to enhance BBB properties. Upon injection, these phenolic sulfates rapidly crossed the BBB, proving their true brain bioavailability.

## Experimental

### (Poly)phenol metabolites chemical synthesis

All LMW (poly)phenol metabolites present in this study were synthesized as before^25^ and characterized by ^1^ H and ^13^C NMR (ESI Fig. S1–S3†). The phenolic pyrogallol-sulfate herein abbreviated as Pyr-sulf, yield 75%, was obtained as a mixture of two isomers of similar proportions (benzene-1,2-diol-3-sulfate and benzene-1,3-diol-2-sulfate, 51 : 49). The phenolic phloroglucinol-sulfate (benzene-1,3-diol-6-sulfate) herein abbreviated as Phlo-sulf, yield 83%, was mainly obtained as mono-sulfate (phloroglucinol-mono-sulfate, phloroglucinol-di-sulfate, phloroglucinol-tri-sulfate, 1 : 0.23 : 0.05, respectively). The phenolic resorcinol-sulfate (phenol-3-sulfate) herein abbreviated as Res-sulf, yield 98%, was obtained as a mixture of resorcinolmono-sulfate (phenol-3-sulfate) and resorcinol-di-sulfate (benzene-1,3-disulfate) (1 : 0.40). All phenolic sulfates were purified by liquid–liquid extraction and liquid chromatography, as described before.^25^ Synthesized compounds were dissolved in DMSO at 4 mM stock concentration before dilution to final concentration in specific cell media.

### Cell culture conditions

HBMEC were routinely cultured in RPMI 1640 medium (Sigma-Aldrich, St Louis, MO, USA) supplemented with 10% fetal bovine serum (FBS), 1% non-essential amino acids, 1% minimal essential medium vitamins, 1 mM sodium pyruvate, 2 mM L-glutamine (Biochrom AG, Berlin, Germany), 10% NuSerum IV (BD Biosciences, Erembodegem, Belgium), and 1% antibiotic-antimycotic solution (Sigma-Aldrich, St Louis, MO, USA).^12^ Cells were maintained in a humidified atmosphere at 37 °C with 5% CO2.

### In vitro BBB transport of (poly)phenol metabolites

For BBB transport evaluation, HBMEC were plated onto rat-tail collagen-I (BD Biosciences) coated semi-permeable membranes of polyester transwell inserts (1.12 cm2, 0.4 μm, Corning Costar Corp., New York, NY, USA), to mimic the blood and brain compartments, at a density of 8 × 104 cells per insert, placed in 12-well plates.^12^ HBMEC were treated after monolayer formation (i.e., after 4 days in culture) and transport assays conducted as previously.12 HBMEC were apically incubated with each metabolite at 5 µM for 15 s, and 2, 5, 15, 30, 120 min (2 h), and medium from both upper and lower compartments collected and immediately frozen at −80 °C until analysis. Monolayer integrity was ensured by the measurement of transendothelial electrical resistance (TEER) and by evaluating alterations in paracellular permeability to sodium fluorescein, as before.^12^

### BBB fluorescence microscopy

For immunofluorescence, HBMEC were seeded onto rat-tail collagen-I coated coverslips at a density of 5 × 104 cells per well in 24-well plates and incubated with each of the (poly)phenol metabolites at 5 μM for 2 h, as before.^26^ HBMEC were incubated overnight at 4 °C with the primary antibodies (antiβ-catenin, #71-2700; anti-ZO-1, #61-7300; anti-CAT1, #PA5-82800, Thermo Fisher Scientific, Waltham, MA, USA; antiASCT1, ab204348; anti-OATP1A2, ab221804, Abcam, Cambridge, United Kingdom; anti-caveolin-1, #3238S, Cell Signaling Technology, Danvers, Massachusetts, USA; anti-Tf (clone HTF-14), #10-258-C025, Enzo Life Sciences, Farmingdale, New York, USA) and thereafter with the corresponding secondary antibodies (Alexa Fluor 594 goat anti-rabbit IgG (H + L), #A11037; Alexa Fluor 488 goat anti-mouse IgG (H + L) #A11001, Thermo Fisher Scientific) for 60 min at room temperature, in the dark. Cells were then mounted on microscopy slides with Prolong Gold Antifade with DAPI (Thermo Fisher Scientific), properly dried, and stored at 4 °C until image acquisition. Between incubations, cells were washed three times with PBS.

### Image acquisition and analysis

Immunolabellings were examined using a Zeiss Z2 microscope equipped with a high-resolution camera (Axiocam 506), using a 40× objective, and equipped with ZEN Blue 2012 software. Ten fields of each condition were acquired and evaluated. Membrane, nuclei, and cell beta(β)-catenin mean fluorescence intensity was quantified using five cells per image in which the “Area”, “Ellipse”, and “Polygon” tools of Icy (Institute Pasteur and France BioImaging, Paris, France) software were used, respectively. The plot profiles and surface plots for beta(β)-catenin protein localization along the cell were obtained by the “Plot Profile” and “Surface Plot” tools in ImageJ (National Institutes of Health, Bethesda, MD, USA) software, respectively. ZO-1 membrane gaps number per cell were quantified in five cells per image, using ImageJ software. Cav1 mean intensity per cell was quantified using five cells per image in which the “Polygon” tool of Icy software was used. The number of Cav1 positive caveolae were quantified using “Spot Detector” tool from Icy software, considering scale 2, sensitivity at 70 and ∼3 pixels size parameter, in five cells per image. Transferrin (Tf) and ASCT1, OATP1A2, CAT1 transporters mean intensity per cell were quantified using ImageJ software, measuring the total image mean intensity and dividing by the number of cells per image.

### Real-time quantitative PCR

Different SLC real-time quantitative PCR analysis (qRT-PCR) was performed in HBMEC, as before.^12^ Oligonucleotide primers were used at a final concentration of 5 μM (ESI Table S1†). Cycles threshold (Ct’s) and melting curves were determined using Thermo Fisher Connect Platform (Thermo Fisher Scientific) (https://www.thermofisher.com/us/en/home/ digital-science/thermo-fisher-connect.html) and results were processed using the 2^−ΔCt^ method for relative gene expression analysis,^12^ and normalized using house-keeping gene β2-microglobulin (B2M) and β-actin.

### In vivo biodistribution and brain uptake of (poly)phenol metabolites

#### Animal treatment

Thirty-six male rats (RccHan®:WIST rat, Envigo RMS S.r.l., Udine, Italy) were housed in the animal facility at the University of Trieste. 12-week-old rats weighing 288 ± 20 g were maintained in temperature-controlled rooms at 22–24 °C, 50–60% humidity, and 12 h light/dark cycles. Animals were fasted overnight before injection. Rats were anesthetized with intraperitoneal administration of Zoletil (25 mg mL−1) and Xylazine (10 mg mL−1). The penis of the rats was extruded by sliding the prepuce downwards. Each rat was i.v. injected with 300 µL PBS containing 50 µL of methanol (vehicle, 3 blank rats) or one of the metabolites (Pyr-sulf, Phlosulf and Res-sulf) single-injected (1 µmol) in 5 Wistar rats per time point into the dorsal penis vein. Rats were sacrificed by decapitation 15 s or 5 min after injection. Blood was drained from the trunk upon head decapitation and urine was collected from the urinary bladder with a syringe. 5 mL of blood were transferred directly to 45 mL of ice-cold methanol : water (95 : 5, v/v), and urine was transferred to 3 mL of ice-cold methanol : water (95 : 5, v/v). The liver, kidney and brain were excised, weighted, washed, immediately frozen in liquid nitrogen and stored at −80 °C. This study has received bioethical approval from the Italian Government (Project Code 4132PAS18, Authorization n.116/2019-PR, by the Ministry of Health). The methods and results comply with the ARRIVE guidelines.^27^

#### Sample preparation

Each frozen tissue was ground under cryogenic conditions using liquid nitrogen with a Cryo Mill and treated as described before.^28,29^ For blood and urine, an equal volume of sample and phosphoric acid (4%) was mixed followed by centrifugation at 8800g for 10 min. Supernatants were pre-treated by Solid Phase Extraction (OASIS HLB 3cc (60 mg) Waters, Milford, MA, USA) to clean up and pre-concentrate the sample,^28,29^ before UHPLC-MS/MS analysis.

### Targeted UHPLC-MS/MS

Both samples from in vitro and in vivo experiments were analyzed by targeted UHPLC-MS/MS. HBMEC medium samples were filtered with a 0.22 μm filter and transferred into liquid chromatography vials before injection. An ExionLC system interfaced with AB6500+ QTrap mass spectrometer with electrospray ionisation system (ESI) (Applied Biosystems/MDS Sciex, Toronto, ON, Canada) was used. All samples were analyzed on a reversed phase (RP) ACQUITY UPLC 1.8 m 2.1 × 100 mm HSS T3 column (Waters) protected with an Acquity UPLC® BEH HSS T3 1.8 m 2.1 × 5 mm precolumn (Waters), at 40 °C and with a mobile phase flow rate of 0.40 mL min−1. Water was used as the weak eluting solvent (A) and acetonitrile as the strong eluting solvent (B); formic acid 0.1% (v/v) was added in both eluents. The multistep linear gradient used was as follows: 0 min 95% A; 0–3 min, 95–80% A; 3–7 min, 80–25% A; 7–8 min, 25–0% A; 8–10.8 min, 0% A isocratic; 10.8–11 min, 0–95% A; 11–13 min 95% A isocratic. Injection volume was 1 µL. The transitions and spectrometric parameters were optimized individually for each standard by the direct infusion of their solutions (10 µg mL−1). The two most abundant fragments, to be used as the quantifier and qualifier, were identified for each compound. Declustering potential (DP) and entrance potential (EP) were optimized for each precursor ion and collision energy (CE) and Collision CellExitPotential (CXP) for each production. The presence of each metabolite of interest was confirmed using the q/Q ratio. The spray voltage was set at −4500 V for negative mode. The source temperature was set at 500 °C, the nebulizer gas (Gas1) and heater gas (Gas2) at 55 and 65 psi, respectively. The spectra derived from the precursor ion’s fragmentation were acquired in trap (enhanced product ion) to confirm the compound with a collision energy of 35 V with CES of 15 V. Calibration curves with in-house synthesized LMW (poly)phenol metabolites were prepared in solvent using 15 levels of concentrations between 0.002 µg L−1 and 10 mg L−1. BBB transport percentage (%) in HBMEC was calculated as before,12 determined by the ratio of lower compartment concentration and the sum of upper and lower compartments concentrations. The permeability coefficient for each metabolite (Papp, cm s−1) was calculated as previously reported.30 Considering both the concentration of metabolites present in the blood and the residual blood volume in the brain, a correction was made to minimize the influence of circulating metabolites on the brain concentration of the metabolites: assuming the blood volume of the rat brain 47.7 μL g−1, 31 the estimated amount of metabolites in the intravascular blood present in the brain was subtracted relative to the total concentration found in the brain.

### Statistical analysis

Results were analyzed using GraphPad Prism® 8.0.2 (GraphPad Software, San Diego, CA, USA). Data represent the average of at least three independent experiments (n = 3). Oneway ANOVA with Dunnet’s test was performed for multiple comparisons between conditions. Comparisons between two groups were performed using two-tailed Student’s t-tests. Data are presented as mean ± SD, or within 25^th^ and 75^th^ percentiles (for RT-qPCR data). Statistically significant differences were considered when p < 0.05.

## Results

### (Poly)phenol metabolites are differently transported across the BBB over time

Despite the acknowledged brain benefits of (poly)phenols from diet, circulating bioavailable (poly)phenol metabolites BBB transport are still marginally understood. With the aim to assess the permeability of the brain microvascular endothelium to three LMW (poly)phenol metabolites detected in human blood, namely Pyr-sulf, Phlo-sulf, and Res-sulf, each metabolite was tested in a cell model of the BBB (Fig. 1) in a transwell system.^12,23^ Each (poly)phenol metabolite was apically applied at physiologically relevant concentrations (i.e., 5 μM), while samples from both apical and basolateral compartments were collected at different timepoints (15 s, and 2, 5, 15, 30, 120 min) for UHPLC-MS/MS analysis (Fig. 1A). Additionally, percentages of transport increased over time, with Pyr-sulf presenting a significantly higher percentage of BBB transport than Phlo-sulf or Res-sulf at 15 min (p < 0.05) and at 30 min (p < 0.001 and p < 0.05, Fig. 1E). Similarly, the significant differences observed in BBB transport percentage were mirrored by an increased BBB permeability of Pyr-sulf comparatively to Phlo-sulf and to Res-sulf at 30 min (p < 0.01 and p < 0.001, respectively, Fig. 1F). None of these significant differences in terms of BBB transport percentage or permeability were reflected in terms of variations in TEER or increased paracellular permeability to sodium fluorescein (Fig. 1G), suggesting that the results obtained do not arise from alterations in monolayer integrity.

**Fig. 1.**
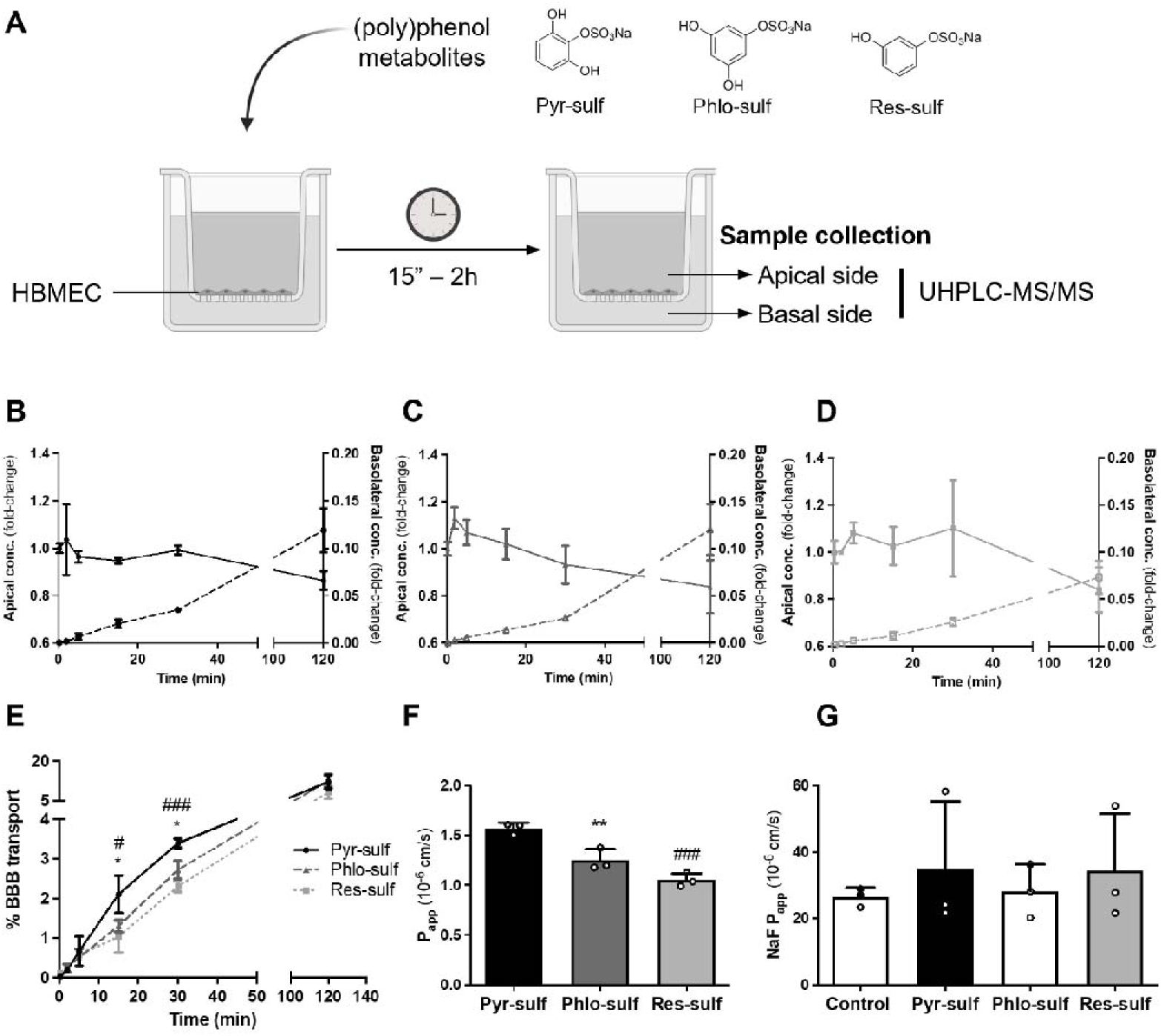
(Poly)phenol metabolites are transported across the blood–brain barrier (BBB) endothelium. (A) Scheme of the experimental design. Human brain microvascular endothelial cells (HBMEC) were grown to confluence on a transwell system. Each of the (poly)phenol metabolites, Pyr-sulf, black, Phlo-sulf, dark grey, or Res-sulf, light grey were apically added at circulating concentrations (5 μM) and samples were collected from both apical and basolateral side at different timepoints (15 s to 2 h) to be analyzed by LC-MS/MS. (B–D) Kinetics of metabolite transport across the BBB endothelium. Apical and basolateral concentrations (conc.) of (B) Pyr-sulf, (C) Phlo-sulf, and (D) Res-sulf are presented in fold-change respectively to time 0. (E) Percentage (%) of BBB transport for each metabolite was calculated from apical and basolateral conc. variations along time. (F) Apparent permeability (Papp) values for each metabolite were obtained for 30 min. (G) Sodium fluorescein (NaF) permeability was monitored to ascertain potential alterations in paracellular transport. Data are given as mean ± SD (n = 3). One-way ANOVA was used to evaluate the significant differences between conditions, presented as *p < 0.05 or **p < 0.01 for Pyr-sulf vs. Phlo-sulf, and as #p < 0.05 and ###p < 0.001 for Pyr-sulf vs. Res-sulf.

### (Poly)phenol metabolites improve barrier features at physiological concentrations

Knowing the three (poly)phenol metabolites transport kinetics across the endothelial cells of the BBB, and bearing in mind that at least one of them, Pyr-sulf, presented pleiotropic potential against oxidative stress, glutamate excitotoxicity, neuroinflammation and dopaminergic cell death,^12,22^ we wondered what their contribution could be to improve barrier tightness. To this end, HBMEC were incubated with each of the metabolites at 5 μM for 2 h, and the expression of both β-catenin and ZO-1 were assessed by fluorescence microscopy (Fig. 2), key proteins for adherens junctions (AJs) and tight junctions (TJs), respectively. We observed that 2 h of incubation was sufficient to rise the expression of β-catenin at cell junctions, reflecting an apparent increase in monolayer organization (Fig. 2A). Semi-quantitative analysis corroborated such observations: Pyr-sulf and Res-sulf led to a significant increase in β-catenin mean intensity per cell (p < 0.001, Fig. 2B), while Pyr-sulf also significantly increased membrane β-catenin (p < 0.001, Fig. 2C). Plot profile and surface plot representations of β-catenin expression on individual cells for each condition reinforce such results, pointing to an enhanced sorting of β-catenin towards the cell membrane, being more evident with Pyr-sulf and Res-sulf (Fig. 2D and E). Regarding the TJs protein ZO-1, we also observed that the incubation with the metabolites led to alterations in this junctional protein organization at cell contacts (Fig. 2F). Such increased monolayer organization could be visible by a clear decrease in the number of ZO-1 gaps per cell (Fig. 2G), which was corroborated by the semi-quantitative analysis performed, highlighting that both Pyr-sulf and Res-sulf significantly reduced the number of ZO-1 gaps per cell (p < 0.001 and p < 0.01, respectively) (Fig. 2H). Collectively, we observed that both Pyr-sulf and Res-sulf can significantly improve BBB tightness properties, with Pyr-sulf being the most promising (poly)phenol metabolite regarding BBB transport and junctional proteins modulation.

**Fig. 2.**
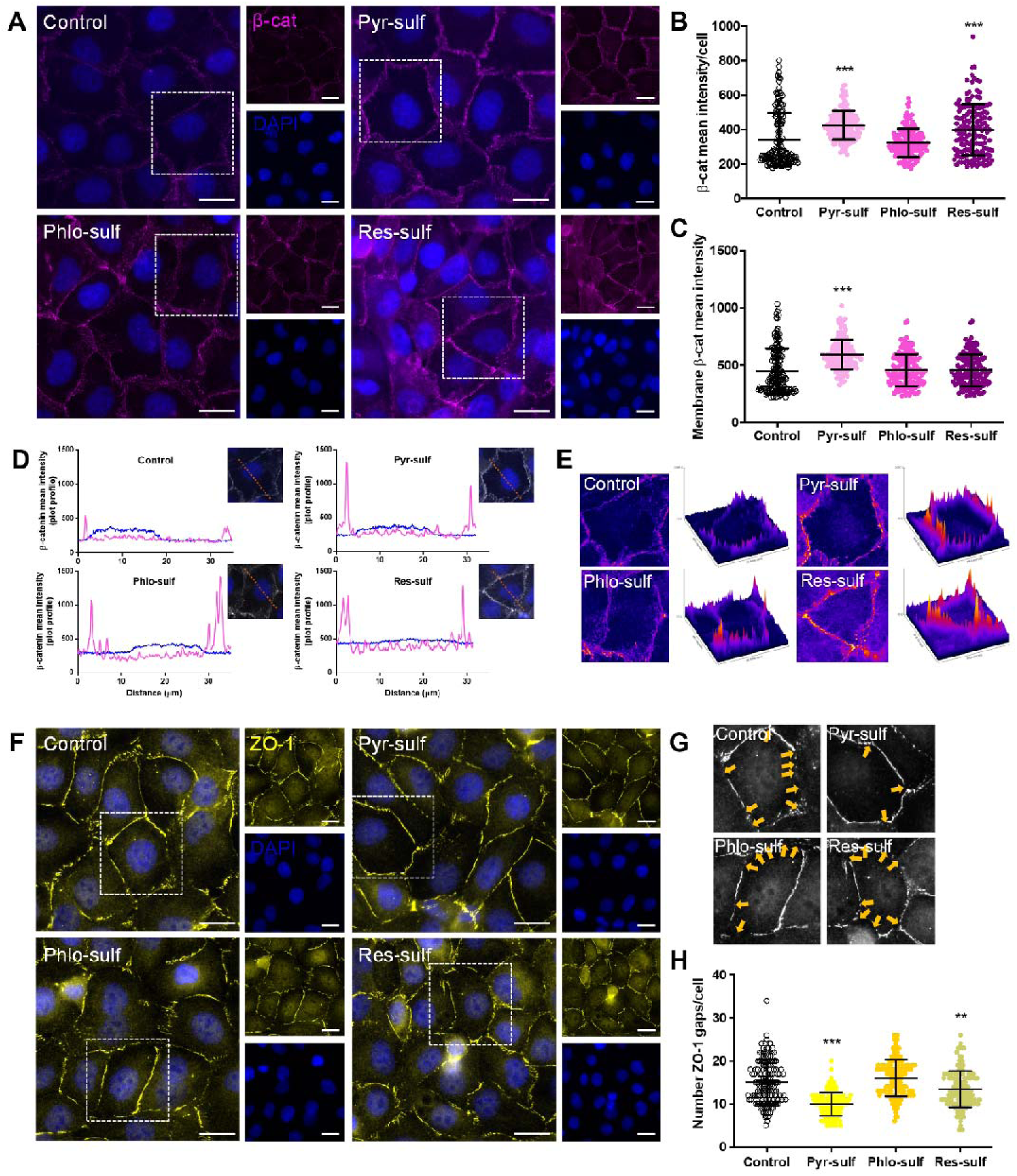
(Poly)phenol metabolites can improve blood–brain barrier (BBB) features. Human brain microvascular endothelial cell (HBMEC) line was used as an in vitro model of the BBB. Each of the (poly)phenol metabolites, Pyr-sulf, Phlo-sulf, or Res-sulf were added to confluent HBMEC monolayers in coverslips at circulating concentrations (5 μM) for 2 h and the expression of adherens and tight junction proteins, β-catenin (β-cat) and zonula occludens-1 (ZO-1), respectively, evaluated by immunofluorescence analysis. (A) Representative images of β-cat (magenta) expression in the different conditions, where white squares identify the cells selected for plot profile and surface plot analysis. Semi-quantitative analysis of (B) mean intensity per cell and (C) on the membrane highlight significant increase of β-Cat by the metabolites, mirrored in the obtained (D) plot profiles and (E) cell surface plots. (F) Representative images of ZO-1 (yellow) expression in the different conditions, where white squares indicate further magnified cells for (G) membrane gap identification (yellow arrows). Semi-quantitative analysis of (H) the number of membrane gaps per cell highlights a significant reduction by Pyr-sulf and Res-sulf. Nuclei counterstained with DAPI (blue). Scale bar: 20 µm. Data are given as mean ± SD (n = 3, 5 cells per image, 10 fields per condition). Oneway ANOVA was used to evaluate the significant differences between conditions, presented as **p < 0.01 and as ***p < 0.001 comparatively to control.

### (Poly)phenol metabolites increase caveolin-1 expression

The mechanisms by which LMW (poly)phenol metabolites can cross the BBB are still quite underexplored. Besides lipophilic or paracellular BBB transport, also vesicular transcytosis may be involved. To shed light on this, we investigated Cav1 protein, which is involved in the formation of flask-shaped invaginations in the plasma membrane mediating transcytosis processes.^32^ Moreover, the possible influence of Tf, involved in endocytosis and recycling mechanisms, regulating iron uptake in almost all cell types,^33^ in favoring the passage of LMW (poly) phenol metabolites across BBB was also investigated (Fig. 3). We observed that 2 h of incubation with each of the metabolites did not induce changes in Tf expression in HBMEC (Fig. 3A). Semi-quantitative analysis showed no significant differences in Tf mean intensity per cell between the different conditions (Fig. 3B), supporting our observations. Regarding Cav1, a clear increase in its expression was observed in HBMEC upon LMW (poly)phenol metabolites incubation (Fig. 3C). Quantification of the fluorescence intensity indicates that Pyr-sulf and Res-sulf exposure led to a significant increase in Cav1 mean intensity per cell (p < 0.001, Fig. 3D). Moreover, spot detector analysis (Fig. 3F) revealed an increase in the number of caveolae per cell upon exposure to Pyr-sulf (p < 0.01), Res-sulf (p < 0.01) and Phlo-sulf (p < 0.05), as documented in Fig. 3E, suggesting an enhanced caveolae-mediated transport, which may comprise one of the pathways responsible by LMW (poly)phenol metabolites transport across the BBB. These data support the view that these phenolic sulfates are transported across the BBB endothelium, possibly aided by transcytosis mechanism mediated by caveolae, rather than by diffusion or paracellular ones.

**Fig. 3.**
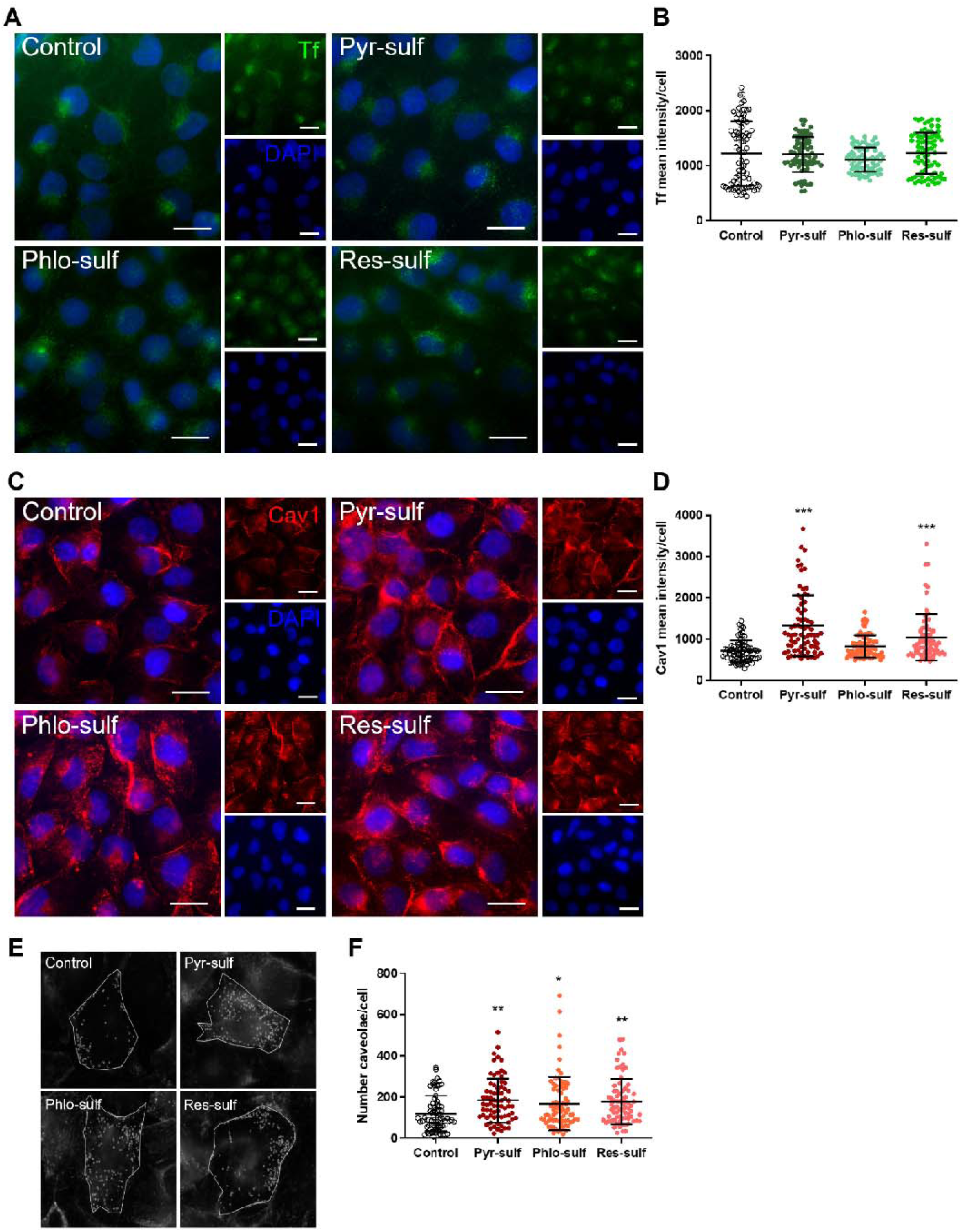
(Poly)phenol metabolites increase caveolin-1 expression but not transferrin. Each of the (poly)phenol metabolites, Pyr-sulf, Phlo-sulf, or Res-sulf were added to confluent HBMEC monolayers in coverslips at circulating concentrations (5 µM) for 2 h and the expression of transferrin (Tf) and caveolin1 (Cav1) was evaluated by immunofluorescence analysis. (A) Representative images of Tf (green) expression in the different conditions. (B) Semi-quantitative analysis of mean intensity per cell. (C) Representative images of Cav1 (red) expression in the different conditions and (D) corresponding semi-quantitative analysis of mean intensity per cell. (E) Representative images of spot detector analysis of Cav1 signal, indicative of (F) total number of caveolae per cell. Nuclei counterstained with DAPI (blue). Scale bar: 20 µm. Data are given as mean ± SD (n = 3; 5 cells per image; 10 fields per condition). One-way ANOVA was used to evaluate the significant differences between conditions, presented as *p < 0.05, **p < 0.01 and as ***p < 0.001 comparatively to control.

### Pyr-sulf modulates the expression of SLC BBB transporters

Since exposure of HBMEC to the metabolites for 2 h proved to significantly modulate the expression of AJs and TJs and increased Cav1 expression and caveolae number, we wondered if specific BBB transporters could also be altered by the most transported (poly)phenol metabolite, Pyr-sulf (Fig. 4), and, in the end, contribute to the BBB permeability observed. We performed a literature revision of the most representative carrier-mediated (CMT) and receptor-mediated transporters (RMT) predicted to be present in brain microvascular endothelial cells^34–38^ (Fig. 4A). To assure which major BBB transporters are expressed in the HBMEC line, a RT-qPCR was performed. Excitatory amino acid transporter 2 (EAAT2/SLC1A2), organic anion transporter 3 (OAT3/SLC22A8), and Ral binding protein (RLIP76/RALPBP1) (an ATP-dependent transporter of glutathione conjugates^39^) appeared as the top three most abundant/ expressed transporters in HBMEC. On the other hand, the SLC21A family transporters (i.e., the organic anion transporting polypeptide 1A2 (OATP1A2/SLCO1A2), and 2B1 (OATP2B1/ SLCO2B1)) together with excitatory amino acid transporter 1 (EAAT1/SLC1A3) and low-density lipoprotein receptor-related protein 1 (LRP1/LRP1) were the ones with the lowest expression levels in HBMEC, representing the least abundant genes (Fig. 4B). For the 25 transporters, we observed that solely 2 h incubation with Pyr-sulf were sufficient to lead to a significant alterations in HBMEC: to our surprise, we observed a downregulation of four CMT, namely the alanine/serine/cysteine/ threonine-preferring transporter 1 (ASCT1/SLC1A4, p < 0.001, Fig. 4C), the cationic amino acid transporter 1 (CAT1/SLC7A1, p < 0.01, Fig. 4D), OATP1A2/SLCO1A2 (p < 0.01, Fig. 4E) and sodium-dependent neutral amino acid transporter-2 (SNAT2/ SLC38A2, p < 0.05, ESI Fig. S4†), not altering the remaining eighteen CMT and none of the three RMT assessed (ESI Fig. S5†). To support our RT-qPCR findings, we also looked for the protein expression by immunofluorescence. Despite no major differences regarding subcellular localization were observed between control and treated HBMEC, Pyr-Sulf was able to significantly reduce the cell immunoreactivity of ASCT1 (p < 0.05, Fig. 4F), CAT1 (p < 0.01, Fig. 4G) and OATP1A2 (p < 0.01, Fig. 4H), possibly mimicking a mild stressor to cells. To the best of our knowledge, this is the first study that elucidates about the relative abundance of the different CMT and RMT in this HBMEC line, and their decrease upon LMW (poly)phenol metabolite exposure in physiological time and concentration.

**Fig. 4.**
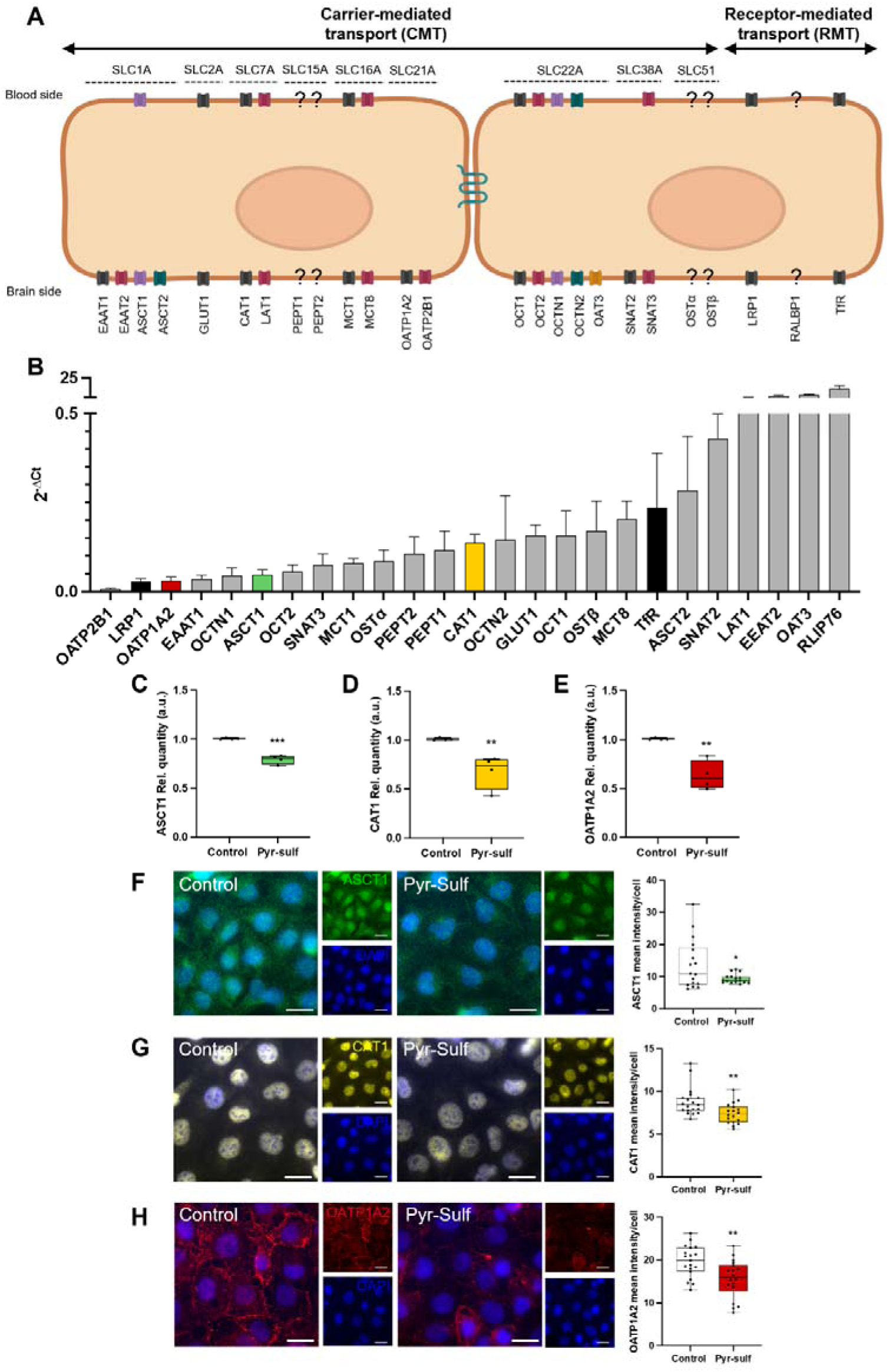
Pyr-Sulf can modulate the expression of carrier-mediated transporter (CMT) genes in human brain microvascular endothelial cells (HBMEC). (A) Schematic representation of CMT and receptor-mediated transporters (RMT) predicted expression/localization in endothelial cells of the BBB based on literature revision.34– 37 Question marks (?) represent still unknown localization of the transporters in apical or basolateral cell membrane. HBMEC untreated and challenged with Pyr-sulf (5 µM) were collected at 2 h and expression of CMT and RMT were analyzed by RT-qPCR and by immunofluorescence analysis. (B) CMT (grey bars) and RMT (black bars) genes abundance represented in relative quantification (2^−ΔCt^) in untreated HBMEC. Pyr-Sulf was able to significant modulate CMT, namely (C) ASCT1 (green), (D) CAT1 (yellow) and (E) OATP1A2 (red). Representative images of ASCT1 (green, F), CAT1 (yellow, G) and OATP1A2 (red, H) are depicted, as well as corresponding semi-quantitative analysis of mean intensity per cell. Nuclei counterstained with DAPI (blue). Scale bar: 20 μm. Relative quantification data are given in Min to Max (n ≥ 3). Housekeeping genes B2M and β-actin mean were used to normalize the relative expression levels. Unpaired t-test was used to evaluate the significant differences between conditions, presented as *p < 0.05, **p < 0.01 and as ***p < 0.001 for Pyr-sulf vs. control. Transporters abbreviations used (protein/gene): Alanine/Serine/Cysteine-preferring Transporter 2 (ASCT2/SLC1A5), Alanine/Serine/Cysteine/Threonine-preferring Transporter 1 (ASCT1/SLC1A4), Cationic amino acid transporter 1 (CAT1/SLC7A1), Excitatory Amino Acid Transporter 1 (EAAT1/SLC1A3), Excitatory Amino Acid Transporter 2 (EAAT2/SLC1A2), Glucose transporter type 1 (GLUT1/SLC2A1), L-type/large neutral amino acid transporter 1 (LAT1/SLC7A5), Low density lipoprotein receptor-related protein 1 (LRP1/LRP1), Monocarboxylate transporter-1 (MCT1/SLC16A1), Monocarboxylate transporter-8 (MCT8/SLC16A2), Organic anion transporting polypeptide 1A2 (OATP1A2/SLCO1A2), Organic anion transporting polypeptide 2B1 (OATP2B1/SLCO2B1), Organic anion transporter 3 (OAT3/SLC22A8), Organic cation/carnitine transporter 1 (OCTN1/ SLC22A4), Organic cation/carnitine transporter 2 (OCTN2/SLC22A5), Organic cation transporter 1 (OCT1/SLC22A1), Organic cation transporter 2 (OCT2/ SLC22A2), Organic solute transporter subunit alpha (OSTα/SLC51A), Organic Solute Transporter Beta (OSTβ/SLC51B), Peptide Transporter 1 (PEPT1/ SLC15A1), Peptide transporter 2 (PEPT2/SLC15A2), Ral binding protein (RLIP76/RALPBP1), Sodium-coupled neutral amino acid transporter (SNAT3/ SLC38A3), Sodium-dependent neutral amino acid transporter-2 (SNAT2/SLC38A2), Transferrin receptor (TfR/TfR).

### (Poly)phenol metabolites reach the brain in vivo

To validate the appearance of the three LMW (poly)phenol metabolites in the brain in vivo, Pyr-sulf, Phlo-sulf, and Ressulf were i.v. injected through a minimally invasive procedure in Wistar rats and quantified 15 s or 5 min after injection (two timepoints also studied in vitro – Fig. 1) in biofluids (blood and urine), in the brain, and in excretory organs (kidney, liver) (Fig. 5A). The presence of 137 LMW (poly)phenol metabolites, already observed in previous studies^10^ were investigated, even in blank group (saline-injected animals). Pyr-sulf and Res-sulf were found in trace amounts in the liver, kidneys, brain (Table 1), and in the blood as well as, being excreted in the urine in negligible amounts (Table 2). No basal levels of Phlosulf were detected in control animals in any of the studied organs nor in biological fluids (Tables 1 and 2). Immediately following injection (15 s), Res-sulf was detected in the blood at approximately 60 µM, whereas both Pyr-sulf and Phlo-sulf attained about 15 µM. At 5 min, Res-sulf and Pyr-sulf decreased to a similar level (10–16 µM), whereas Phlo-sulf increased to 40 µM (Fig. 5B). The data show that each metabolite was taken into the organs already 15 s after injection. Phlo-sulf presented the highest concentration in both the kidneys (Fig. 5D) and liver (Fig. 5E), as well as in the urine (Fig. 5C) at all times assessed. Of most importance, at 15 s, each metabolite appeared in the brain at concentrations well above their basal values and either increased (Pyr-sulf) or remained stable (Phlo-sulf and Res-sulf) at 5 min, proving that all the metabolites injected in the bloodstream were indeed able to cross the BBB (Fig. 5F). Noteworthily, Res-sulf accumulated in the brain up to 272 times the baseline concentration found in the saline-injected rats at 5 min (Fig. 5F and Table 2). At a distinct extent, Pyr-sulf accumulated in the brain at 0.29 nmol g−1 at 15 s (approximately 9 times more than blank levels) and significantly increased at 5 min to 1.1 nmol g−1 (36-fold change t5/blank) (Fig. 5F and Table 1). Regarding Phlo-sulf, although it was not detected in the brain of control animals, it was found in considerable amounts already 15 s after being injected (∼1.4 nmol g−1) (Fig. 5F and Table 1). As expected, we observed that the three metabolites were excreted in the urine (Fig. 5C) after prior accumulation in the kidneys (Fig. 5D). Collectively, we showed that the three LMW (poly)phenol metabolites studied are present in considerable amounts in the urine and bloodstream, being detected in peripheral organs and reaching the brain in vivo.

**Fig. 5.**
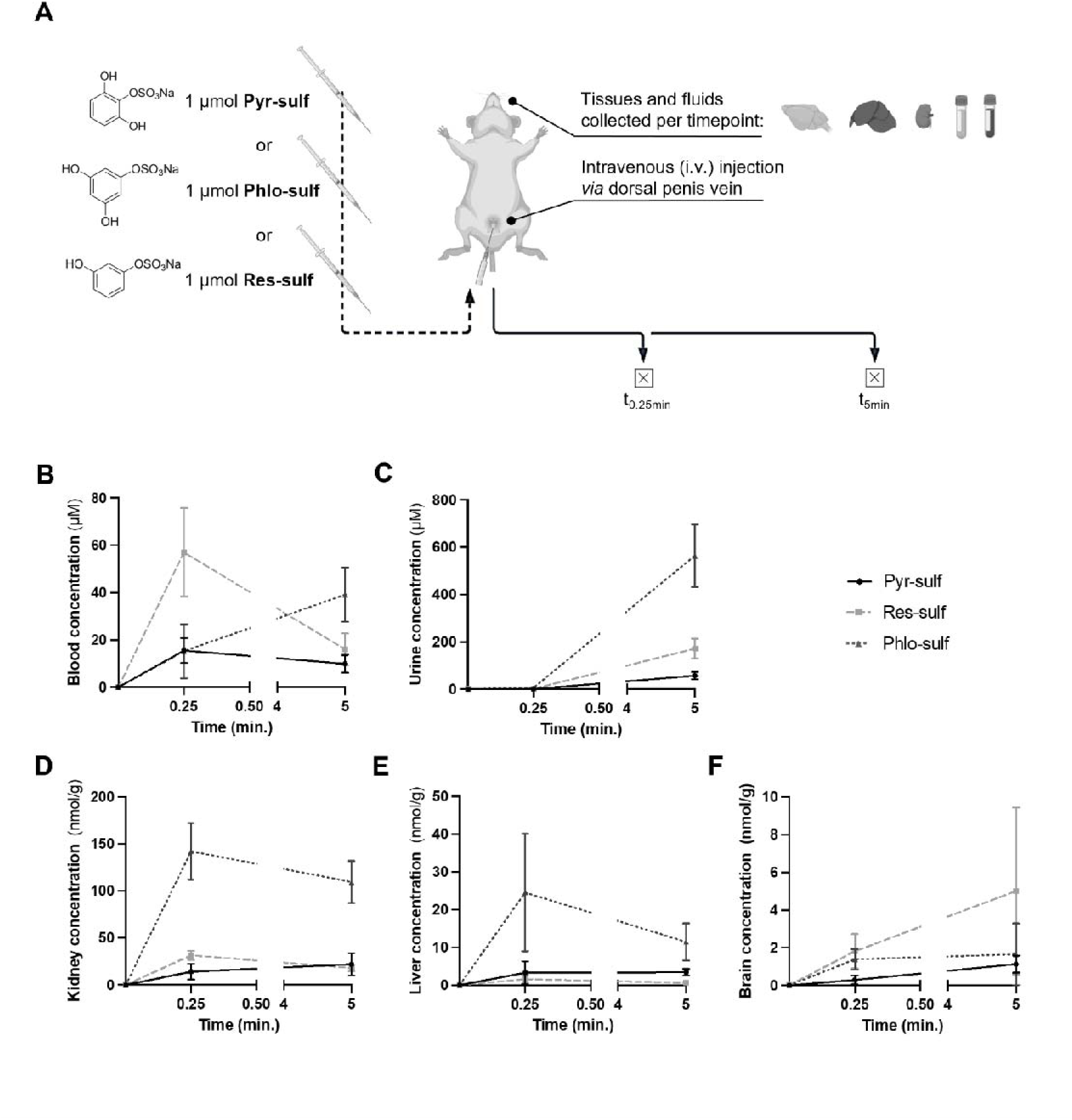
(Poly)phenol metabolites reach the brain upon injection in vivo. (A) Male Wistar rats were intravenously injected (through the dorsal penis vein) with 1 µmol of each of the metabolites and sacrificed 15 s or 5 min after injection. Graphical representation of biofluids, namely (B) blood and (C) urine concentrations of Pyr-sulf, black, Phlo-sulf, dark grey, and Res-sulf, light grey at different timepoints. Graphical representation of (D) kidney, (E) liver, and (F) brain concentrations of each of the metabolites at different timepoints. Data are given as mean ± SD (n = 5).

**Table 1:** Concentrations and percentage (%) of (poly)phenol metabolites recovery in kidney, liver and brain compared to initial injected dose. Pyrsulf, Phlo-sulf, and Res-sulf concentrations measured in the different organs is presented in nmol per gram (nmol g−1) of tissue weight ± SD, where SD is the standard deviation of 5 replicates. For the recovery, the values given are the % of dose found in rats after the i.v. injection at two different time points (t0.25 and t5 min) and control. One-way ANOVA was used to evaluate the significant differences along time between control vs. t0.25 or t5, indicated by the p value, and t0.25 vs. t5, presented as a p < 0.05, b p < 0.01 or c p < 0.001.

|  |  | Kidney |  |  | Liver |  |  | Brain |  |  |
| --- | --- | --- | --- | --- | --- | --- | --- | --- | --- | --- |
| | | nmol/g ± SD | Recovery % | p-value | nmol/g ± SD | Recovery % | p-value | nmol/g ± SD | Recovery % | p-value |
| Pyr-sulf | Blank | 0.209 ± 0.255 | 0.05 | | 0.010 ± 0.005 | 0.01 | | 0.031 ± 0.024 | 0.006 | |
| | t<sub>0.25</sub> | 14.1 ± 8.5 | 3.05 | **0.039** | 3.29 ± 3.06 | 3.04 | **0.023** | 0.29 ± 0.24 | 0.05 | 0.334 |
| | t<sub>5</sub> | 22.0 ± 12.1 | 5.0 | **0.002** | 3.39 ± 0.89 | 2.94 | **0.019** | 1.2 ± 0.5 <sup>b</sup> | 0.2 | **<0.001** |
| Phlo-sulf | Blank | N/D | N/A |  | N/D | N/A |  | N/D | N/A |  |
| | t<sub>0.25</sub> | 142.2 ± 30.2 | 30.9 | **0.003** | 30.6 ± 8.9 | 27.9 | **0.014** | 1.4 ± 0.5 | 0.2 | 0.074 |
| | t<sub>5</sub> | 109.4 ± 22.5 | 22.9 | **0.014** | 11.5 ± 4.9 <sup>a</sup> | 10.09 | 0.343 | 1.7 ± 1.6 | 0.3 | 0.597 |
| Res-sulf | Blank | 0.014 ± 0.011 | 0.003 | | 0.016 ± 0.002 | 0.01 | | 0.018 ± 0.008 | 0.003 | |
| | t<sub>0.25</sub> | 31.5 ± 4.9 | 6.8 | **<0.001** | 1.593 ± 1.002 | 1.44 | **0.044** | 1.8 ± 0.9 | 0.3 | 0.422 |
| | t<sub>5</sub> | 18.1 ± 3.5 <sup>c</sup> | 3.8 | **<0.001** | 0.616 ± 0.404 | 0.53 | 0.431 | 5.0 ± 4.4 | 0.9 | **0.014** |

**Table 2:**
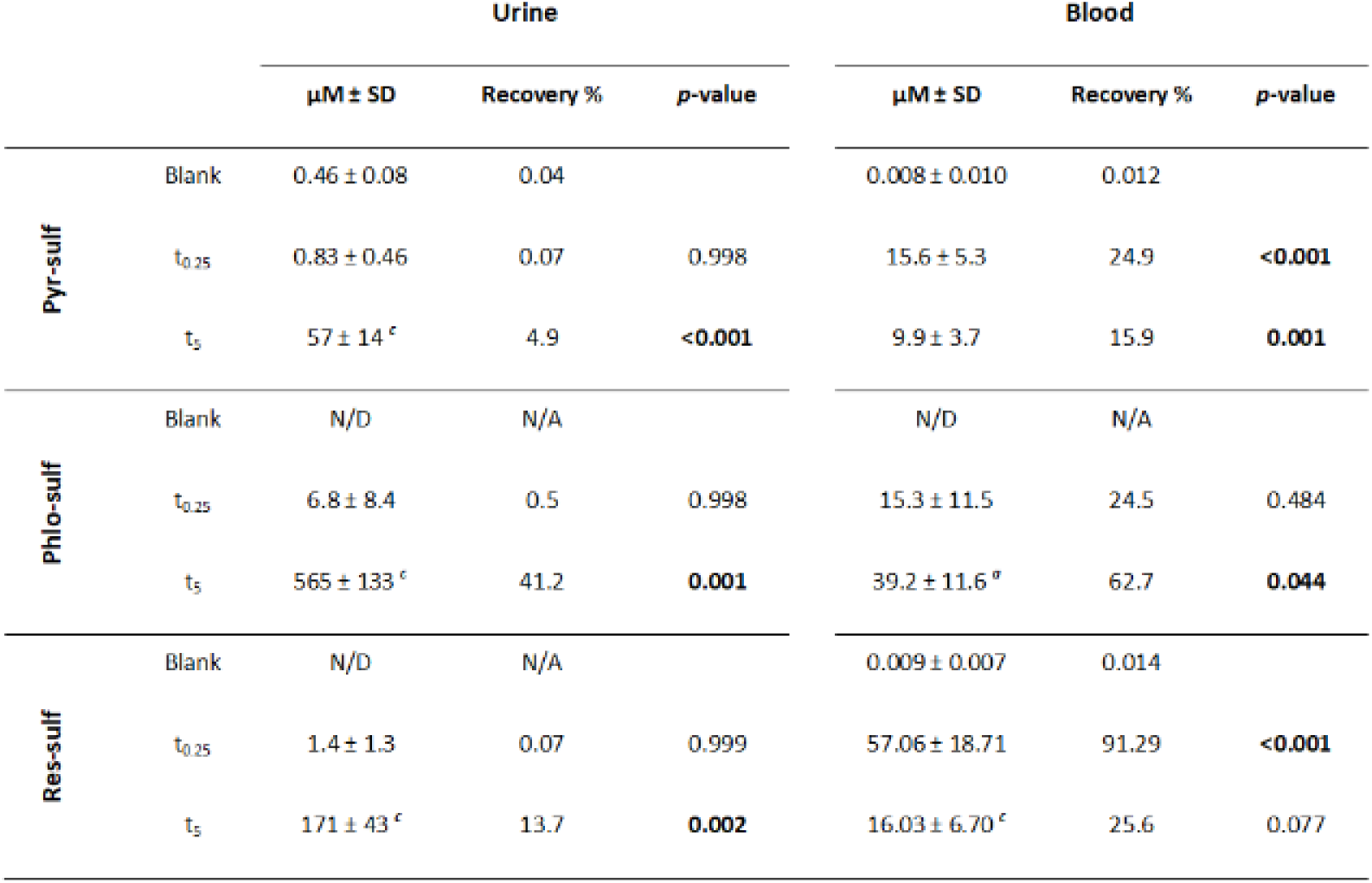
Concentrations and percentage (%) of (poly)phenol metabolites recovery in urine and blood compared to initial injected dose. Pyr-sulf, Phlo-sulf, and Res-sulf concentrations measured in the different biofluids is presented (μM) ± SD, where SD is the standard deviation of 5 replicates. For the recovery, the values given are the % of dose found in rats after the i.v. injection at two different time points (t0.25 and t5 min) and control. One-way ANOVA was used to evaluate the significant differences along time between control vs. t0.25 or t5, indicated by the p value, and t0.25 vs. t5, presented as a p < 0.05, b p < 0.01 or c p < 0.001.

## Discussion

When consumed, phenolic compounds from food can combine with those generated by gut microbiota breakdown of flavonoids and chlorogenic acids, leading to elevated levels of LMW metabolites in circulation.^10^ Among those are phenolic sulfates, like Pyr-sulf, Phlo-sulf, or Res-sulf, found in the bloodstream after the consumption of (poly)phenol-rich diets.^10^ However, our current understanding of the extent to which these metabolites can reach the brain remains limited. Previously, we have reported both isomers of Pyr-sulf BBB endothelial transport and their role attenuating neuroinflammation.^12^ In this study we aimed to further explore Pyr-sulf potential on HBMEC properties, validating its BBB transport in vivo, side-by-side with two structurally similar compounds, Phlo-sulf and Res-sulf. Despite chemically similar, transport kinetics using HBMEC highlighted slightly different LMW (poly)phenol metabolites BBB permeability rates. Theoretically, the absence of one hydroxyl group on Res-sulf comparatively with the other two should improve its BBB transport by passive diffusion, mainly due to increased lipophilicity. Nonetheless, our data points in the opposite direction since Res-sulf presented the lowest endothelial permeability. Indeed, Pyr-sulf emerged as the one with the highest BBB transport in HBMEC, visible already at 30 min. Previous work from our lab using the same BBB in vitro model has shown that Pyr-sulf isomers were transported across the endothelium after 2, 6 and 24 h of incubation,^12^ corroborating our observations for shorter periods of time. Moreover, endothelium permeability for other (poly) phenol metabolites using this and other BBB models have already been reported. It was shown that the metabolite 5-(4′-hydroxyphenyl)-γ-valerolactone-3′-sulfate is transported across HBMEC, being also detected in the brain of rats injected with the aglycone form, 5-(3′,4′-dihydroxyphenyl)-γ-valerolactone.^40^ Hesperetin, naringenin, and in vivo metabolites, as well as dietary anthocyanins and cyanidin-3-rutinoside and pelargonidin-3-glucoside, have also been shown to be uptake by BMEC from mouse and rat.^14^ How exactly the transport occurs at the BBB level, whether it is mainly passive or carrier-mediated, is still not clear. What we observed is that neither the differences in the chemical structure, nor compound lipophilicity are enough to justify the increased Pyr-sulf and Phlo-sulf in vitro BBB transport comparatively to Res-sulf, suggesting that some sort of active transport might be involved. In fact, 3,4,5-trihydroxybenzoic acid (gallic acid), a structurally similar metabolite, has been shown to be a potent inhibitor of OAT1 and OAT3 transporters, where OAT1 was capable of significantly increase gallic acid uptake.^41^ Moreover, 3,4-dihydroxyphenylacetic acid and homovanilic acid (4′-hydroxy-3′-methoxybenzoic acid), two LMW (poly)phenol metabolites and products of dopamine metabolism, have been shown to be transported across the BBB via OAT3.^42,43^ Likewise, we cannot exclude the contribution of some sort of active transport. Aiming to further dissect this possibility, we studied the expression and possible contribution that the most transported metabolite, Pyr-sulf, may have to different SLC transporters. Considering the most documented CMT and RMT at BBB level, we tested 25 transporter candidates for HBMEC.^34–37^ Surprisingly, PyrSulf significantly downregulated four CMT, ASCT1/SLC1A4, CAT1/SLC7A1, OATP1A2/SLCO1A2 at mRNA and protein level and SNAT2/SLC38A2 at mRNA level. Such unexpected alterations may suggest a putative mild-stressor role of Pyr-sulf, decreasing SLC expression in HBMEC, as described to occur in certain pathologies or in response to toxins/drugs.^44^ We can speculate that Pyr-sulf, as previously observed for other (poly) phenols^45^ and by our team in neuronal cells,^22^ may be acting via a pre-conditioning effect as a possible regulation of import protective mechanism, priming HBMEC to better respond to a stronger later insult. Nevertheless, to fully clarify this, HBMEC pre-treatment with Pyr-sulf prior an external insult, as well as the study of each SLC activity in the presence of this LMW should be envisaged. Importantly, the biological role of ASCT1/SLC1A4 in the brain has not been totally elucidated, though its involvement on neutral amino acids traffic across has been reported.^46^ Despite no data available regarding LMW metabolites on SLC, a recent study showed that lemon peel flavonoids containing isomangiferin, rutin, astragalin, naringin, and quercetin, can upregulate mRNA levels of ASCT1/SLC1A4 in response to damage, inflammation, or atrophy of muscle tissue in exhaustive exercised mice.^47^ Similarly to ASCT1/ SLC1A4, the effects of LMW metabolites on the modulation of OATP1A2/SLCO1A2 transporter at BBB level are scarce; what was already reported is that fruit juices may inhibit the absorption of OATP1A2-substrate drugs in the intestine,^48^ while the dietary (poly)phenols apigenin, quercetin and kaempferol have shown to be capable to decrease OATP1A2-mediated transport in kidney cells.^49^ Pyr-sulf downregulation of these SLC transporters at transcriptional and protein levels may indicate a similar mechanism of action, worthwhile to be further explored. What is acknowledged is that, besides direct binding and inhibition, OATPs are regulated at transcriptional and post-translational levels, ultimately controlling the expression of active transporters at the cell surface.^50^ Vesicular transport may also be involved in metabolites BBB transport. Caveolae are specialized lipid rafts, vital for endocytosis and transcytosis processes across the BBB.^51^ The presence of Cav1 is essential in the formation of caveolae, and its increased expression, as induced by the LMW (poly)phenol metabolites, suggests a potential enhancement in caveolarmediated transcytosis. Caveolae importance in transporting nutrients, signaling molecules, and certain drugs across the BBB,^52^ emphasizes the relevance of our observations. Similarly, we have recently observed Cav1 involvement in BBB transport of sesquiterpene lactones from chicory^26^ and it was also reported that salvionic acid, a phenolic compound from Salvia miltiorrhiza, can modulate caveolae-mediated endocytosis at the BBB level, increasing the brain accumulation of doxorubicin.^53^ On the other hand, Tf, which is mainly known for its role in iron transport through receptor-mediated transcytosis, has also been implicated in the transport of other substrates, including drugs and peptides.^54^ The absence of change in its expression in HBMEC upon metabolites treatment might indicate that their BBB transport is not favored by the Tf-receptor-mediated pathway. Nevertheless, further research will be needed to elucidate that. Besides BBB permeability, the fact that they are abundant in circulation makes them candidates not only to act at brain level but before that, at vascular level. Pyr-sulf and Res-sulf presented a strong potential to modulate BBB properties when tested in circulating concentrations, visible in AJs and TJs expression in HBMEC by increasing β-catenin membrane localization and intensity and reducing the number of ZO-1 membrane gaps, respectively, suggestive of increased monolayer integrity and improvement of barrier tightness. Other parent flavonoids (i.e., non-bioavailable metabolites), like epigallocatechin gallate, genistein, myricetin, and quercetin, have already been described to exhibit similar effects on intestinal barrier functions and in the prevention against insults.^55^ At brain level, hesperidin was shown to protect the BBB during ischemia/hypoxia, maintaining the integrity of claudin-5 and ZO-1, both in vitro and in vivo, ^56^ while baicalin increased BBB tightness via claudin-5 and ZO-1.^57^ Also kaempferol attenuated LPS-induced BBB dysfunction by counteracting the increase of BBB permeability and the degradation of TJs proteins such as occludin-1 and claudin-1, as well as the gap junction protein connexin 43.^58^ Regarding metabolites, indole-3-propionic acid was shown to attenuate hypoxicischemic-induced BBB leakage in neonatal rats, holding protective effects on rat BMEC cell viability and proliferation, as well as in AJs and TJs proteins modulation.^59^ In male Wistar ischemic rats exposed to dusty particulate matter, it was observed that pretreatment with gallic acid reversed BBB leakage and lipid peroxidation.^60^ Indeed, gallic and caffeic acid were shown to protect against hyperglycemia-induced BBB disruption in bEnd.3 cells, with caffeic acid increasing claudin-5, occludin, ZO-1 and ZO-2.^61^ Moreover, intraperitoneal injection of protocatechuic acid (3,4-dihydroxybenzoic acid) improved functional recovery after spinal cord injury by attenuating blood-spinal cord barrier disruption and hemorrhage in rats.^62^ Our results with phenolic sulfates, though without any insult, support these observations, with a clear potential already observed after solely 2 h of incubation in HBMEC. This warrants further investigation as a promising strategy for addressing BBB leakage issues or neurological diseases. Nevertheless, as far as we know, this is the first work evidencing the potential of bioavailable phenolic sulfates, when tested at circulating concentrations, to enhance BBB barrier tightness via TJs and AJs modulation in HBMEC. Aiming to validate the true in vivo appearance of these metabolites inside the brain, an animal study was conducted.^24^ We observed that these three phenolic sulfates are almost immediately detected in the brain upon injection, in alignment with what was already reported for other LMW metabolites.^24,30^ Considering the rat blood volume as 64 mL per kg body weight and an average weight of 288 g, the initial concentration for each metabolite in the blood was approximately 62.5 µM, while the global brain concentrations of these metabolites reached the range of 1–2 µM which are appropriate for effective binding and target modulation (enzymes, receptors, transporters, channels).^63^ A potential limitation of our study is the lack of brain perfusion prior to metabolite analysis. The absence of this step might introduce inaccuracies in the measured concentrations of metabolites due to the presence of residual blood in the brain. To address this, we applied a correction by considering both the concentration of metabolites present in the blood and the estimated residual blood volume in the brain. The modest transport percentage of each metabolite in the brain (less than 1%) vs. other tissues reinforces proper BBB function and role to safeguard the brain homeostasis, preventing potentially high harmful concentrations. Moreover, given our previous evidence of further metabolism at the BBB level^12^ this might account for the reduced percentages of BBB transport observed: LMW (poly) phenol metabolites may not only be uptake by HBMEC but also transformed into new end-route metabolites, whose effects are yet to be studied. Apart from Pyr-sulf, the other two metabolites occurred at similar levels in the brain between 15s and 5 min after injection. Unfortunately, we were not able to discriminate the relative abundances of the different isomers of each LMW (poly)phenol metabolite inside animal brains. Nevertheless, we have recently quantified several LMW (poly) phenol metabolites on human CSF samples and we observed that the two isomers of Pyr-sulf concentrations differ among subjects, ranging from 3.8 ± 0.15 nM for pyrogallol-1-sulfate (benzene-1,2-diol-3-sulfate) and 2.1 ± 0.27 nM for pyrogallol-2-sulfate (benzene-1,3-diol-2-sulfate) pointing to pyrogallol-1-sulfate to be almost twice as concentrated than pyrogallol-2-sulfate.^21^ Such observations may already indicate that the brain uptake and bioavailability of different isomers from phenolic metabolites may differ and, therefore, their bioactivity potential may also vary. Nevertheless, all these compiling data support our BBB transport results for both isomers here and before.^12^ Despite the similarities in their chemical structure, the three metabolites presented very distinct profiles in blood at 15 s and 5 min. Pyr-sulf reached a steady state rapidly, while Res-sulf significantly decreased its concentration along time. Phlo-sulf, however, had an unexpected increase in concentration, raising the question of what would happen in longer timepoints. Notwithstanding, as previously reported,^21^ we are aware that, at least for the two Pyr-sulf isomers, a saturation of the concentration was observed in the CSF of human subjects, since an increase in micromolar concentrations in the plasma did not significantly change the nanomolar concentrations in CSF. Such saturation observations for Pyr-sulf can at least partially justify the differences in recovery observed compared with the other catechol metabolites and help to explain Phlo-sulf unexpected increase in concentration. We may speculate that Phlo-sulf may increase in blood with time because it is first rapidly transported into the liver and kidney which in turn decreased its concentration over time. Here, it might be excreted in the bile and urine; however, when efflux transporters pumping it into the excretory compartments are saturated, Phlo-sulf intracellular concentration may increase and promote reflux into the blood. Other interpretation could be that Phlo-sulf may be formed from hydroxylation of Res-sulf at the liver, which then decreases in the blood. Nevertheless, our data do not allow us to estimate the total amount of compound recovery as only some major organs were collected and the metabolites could also have been distributed in larger tissues, like adipose, endothelium, or connective tissue.^24^ Gasperotti et al. likewise share this interpretation: they injected rats with a mixture of 23 gut-microbiota-derived metabolites, which are known to be byproducts of the metabolic breakdown of (poly)phenols, including phloroglucinol (benzene-1,3,6-triol) and pyrogallol (benzene-1,2,3-triol), with the latter never being detected, whereas no sulfate conjugates were present in the mixture.^24^ Our experimental design, which overcomes previous methodological limitations (LC-MS analytic method tailored for the detection and quantification of phenolic sulfates), involving a single injection of each metabolite, might mitigate several potential issues by injecting a multi-component mixture, like the biotransformation of certain metabolites into others that are already present in the mixture and/or potential interactions among metabolites competing for the same enzymes.

## Conclusions

Altogether, our data provides new insights for circulating LMW (poly)phenol metabolites BBB transport. Pyr-sulf was the one that reached the highest BBB transport rate in vitro and the fastest steady state in vivo. Moreover, in physiological concentrations and resident time, Pyr-sulf also downregulated SLC transporters, increased caveolae formation and improved BBB junctional proteins expression, opening doors for future studies as a barrier properties improvement candidate.

## Data availability

All the data for this article, including all the raw data regarding LC-MS, microscopy, RT-qPCR which was used to elaborate the manuscript figures and ESI can be found at: Raw data.†

## Author contributions

Conceptualization, I. F., C. N. S. and S. P.; methodology, R. C., Da. M., D. C., Do. M., M. G.-A., F. T., S. P., and I. F.; formal analysis, R. C., Da. M., D. C., Do. M, M. G.-A., I. F.; investigation, R. C., Da. M, D. C., F. T., S. P., I. F.; writing – original draft preparation, I. F.; writing – review & editing, all; supervision, I. F. and C. N. S.; funding acquisition, S. P. and C. N. S.

## Acknowledgements

To European Research Council (ERC), Grant no. 804229; to iNOVA4Health (LISBOA-01-0145-FEDER-007344; UIDB/04462/ 2020), and iMed (UIDB/04138/2020 and UIPD/04138/2020) cofunded by Fundação para a Ciência e Tecnologia (FCT)/ Ministério da Ciência e do Ensino Superior (MCTES), through national funds, and by FEDER under the PT2020 Partnership Agreement; to FCT via PERCEPT (2022.02127.PTDC) to the European Regional Development Fund through Interreg V-A Italy-Slovenia 2014–2020 (Agrotur II, 1473843258); to the ERDF 2014–2020 Program of the Autonomous Province of Trento (Italy) with EU co-financing (Fruitomics). To FCT for financial support of R. C. (PD/BD/135492/2018), D. C. (2020.04630.BD), D. M. (2021.05505.BD) and I. F. (2022.00151.CEECIND) and to Marco Stebel for his technical support with animal experiments.

