## Supplementary Material for "Circulating low-molecular-weight (poly)phenol metabolites in the brain: unveiling in vitro and in vivo blood–brain barrier transport"

[1] iNOVA4Health, NOVA Medical School, Faculdade de Ciências Médicas, Universidade NOVA de Lisboa, Campo dos Mártires da Pátria, Lisboa, Portugal.

[2] Instituto de Tecnologia Química e Biológica António Xavier, Universidade NOVA de Lisboa, Avenida da República, Oeiras, Portugal.

[3] Metabolomics Unit, Research and Innovation Centre, Fondazione Edmund Mach (FEM), via E. Mach 1, San Michele all'Adige, Italy.

[4] Department of Life Sciences, University of Trieste, via L. Giorgieri 1, Trieste, Italy.

[5] Research Institute for Medicines, Faculty of Pharmacy, Universidade de Lisboa, Av. Prof. Gama Pinto, Lisboa, Portugal.

[6] Department of Pharmaceutical Sciences and Medicines, Faculty of Pharmacy, Universidade de Lisboa, Av. Prof. Gama Pinto, Lisboa, Portugal.

[7] iBET, Instituto de Biologia Experimental e Tecnológica, Avenida da República, Apartado 12, Oeiras, Portugal.

### SUPPLEMENTARY MATERIAL

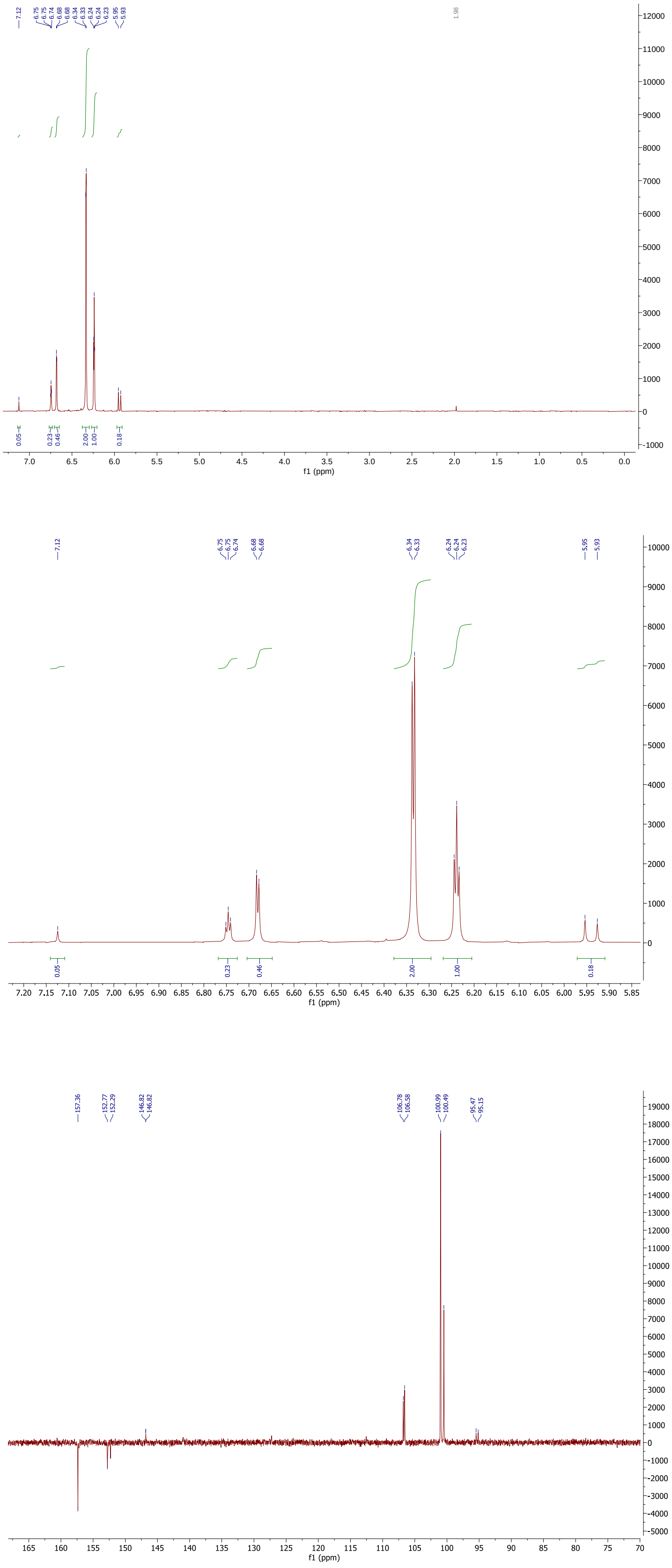


**Fig. S1.** 400MHz NMR spectra of phloroglucinol-*O*-sulfate (**A**) $^{1}$H proton (**B**) $^{13}$C carbon spectra

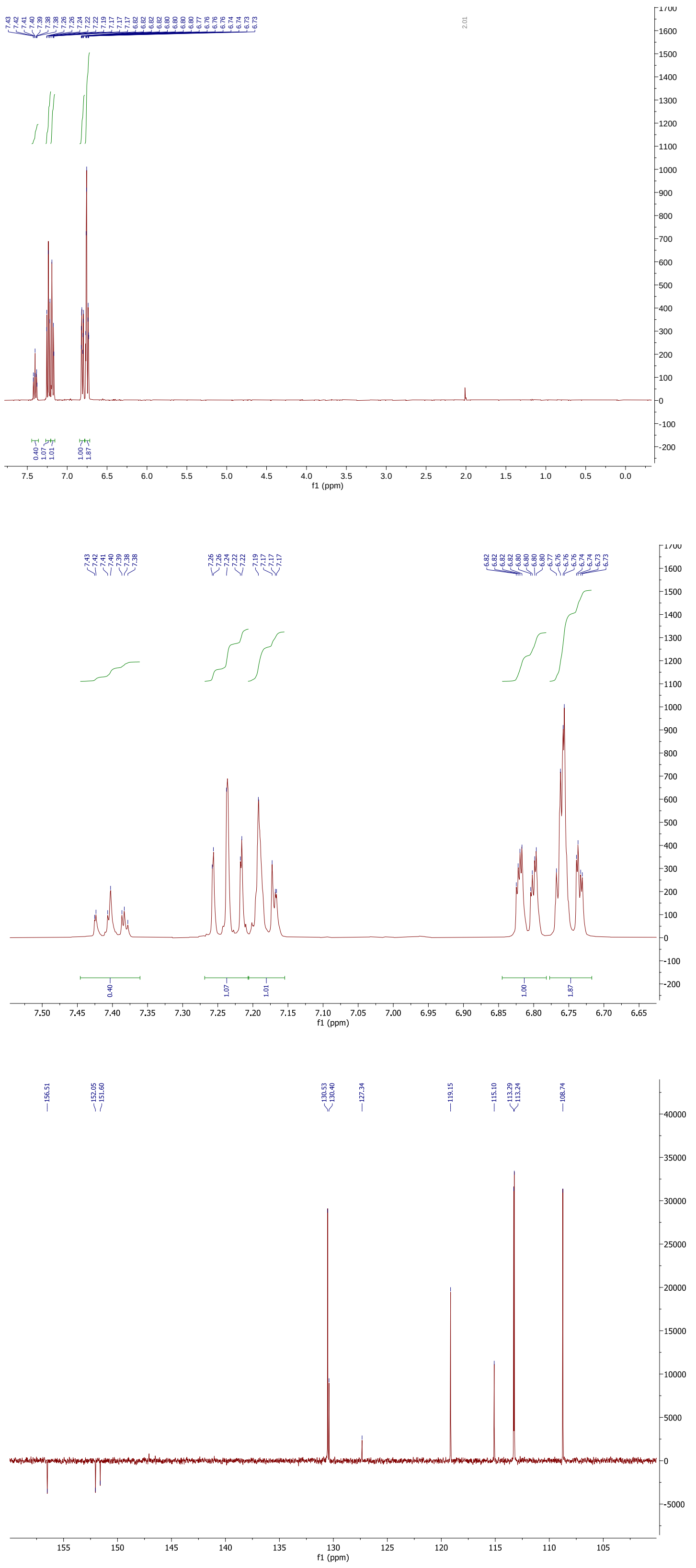

**Fig. S2.** 400MHz NMR spectra of resorcinol-sulfate (A) $^{1}$H proton (B) $^{13}$C carbon spectra

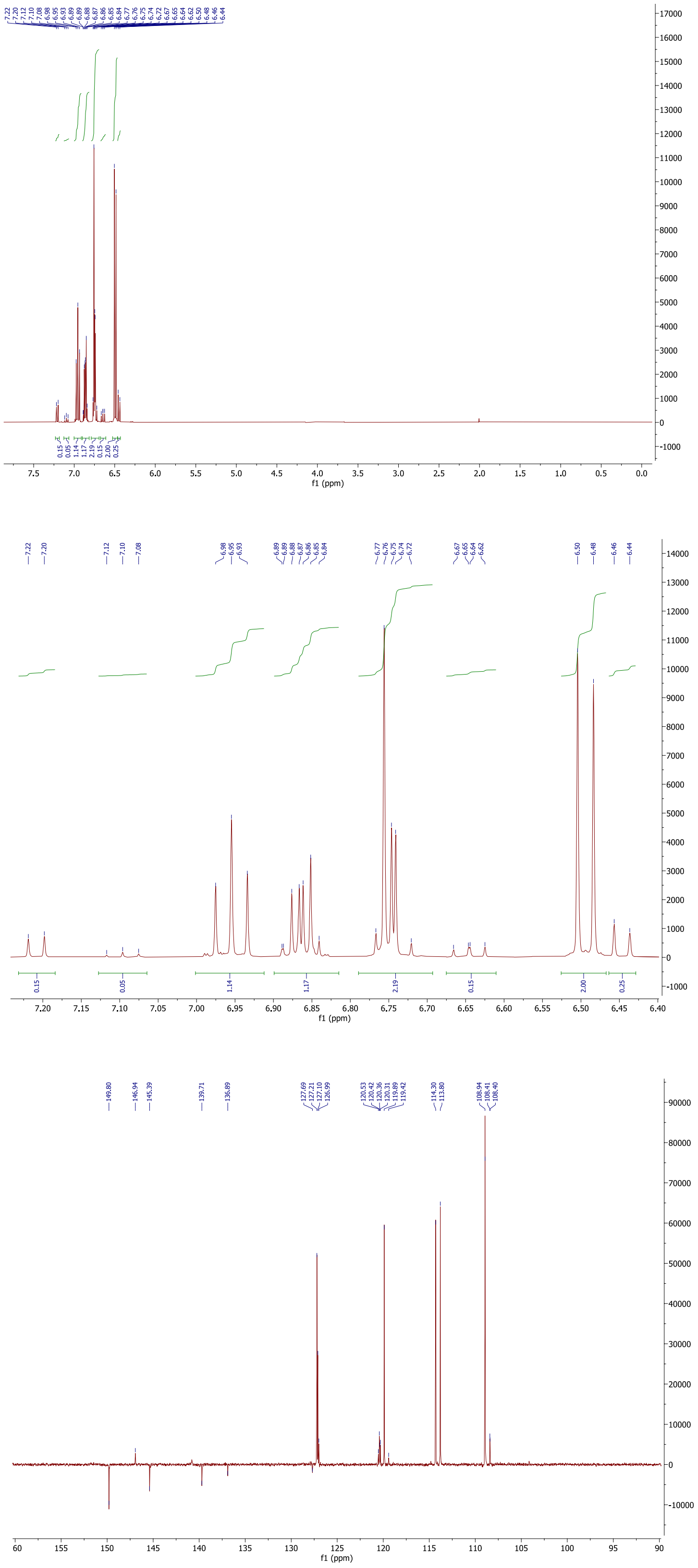


**Fig. S3.** 400MHz NMR spectra of pyrogallol-*O*-sulfate (**A**) $^{1}$H proton (**B**) $^{13}$C carbon spectra

A

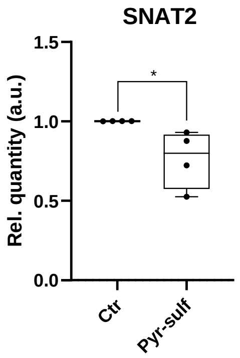


**Fig. S4.** HBMEC incubation with 5μM of Pyr-sulf for 2h significantly modulate the expression of *SNAT2* gene.

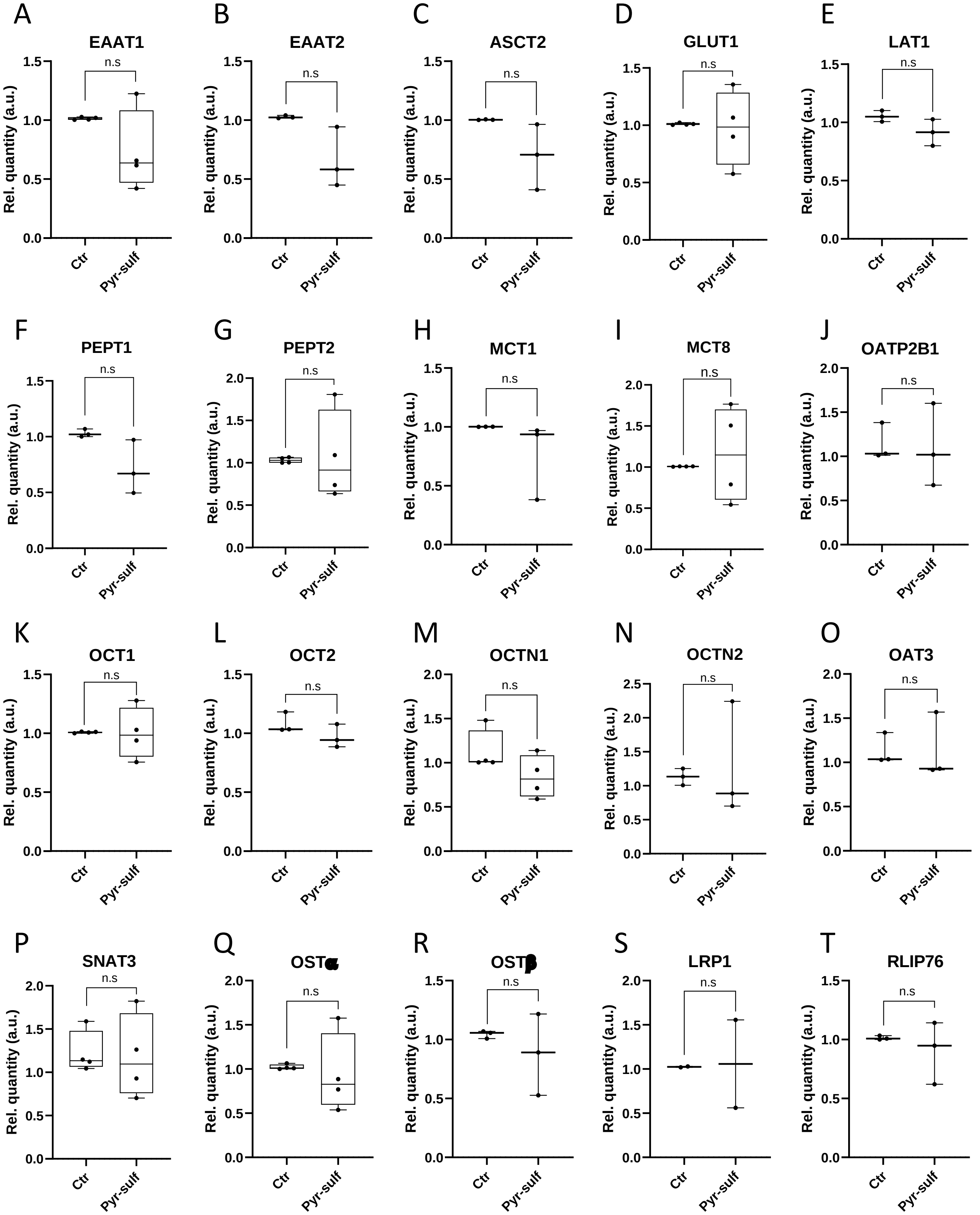


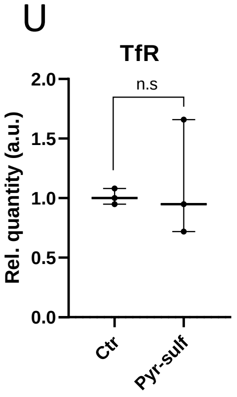


**Fig. S5.** HBMEC incubation with 5µM of Pyr-sulf for 2h do not significantly modulate the expression of SLC transporters.

**Table S1.** Primers list for RT-qPCR analysis of the different human transporters in HBMEC

| Gene (transporter) | Forward (5'→3') | Reverse (5'→3') |
|---|---|---|
| LRP1 | TTGGATTGACGCCAGGTCAG | CCAGTGTGTTTGTTCGCCAG |
| RALPBP1 (RLIP76) | GTAAACAGGCAGAGGCTGGG | AGGGACGAGGTCTTCTCTGT |
| SLC1A2 (EAAT2) | AGAGGGATCGCCTGCAAATC | CAGAGAGTGGTGGCAGAGGA |
| SLC1A3 (EAAT1) | ATTCCAGCAGGGAGTCCGTA | CCAGATGCCCCAGAGAAACAA |
| SLC1A4 (ASCT1) | CTCTCCTCGCCTTTCTCGCA | CTCTCCTCGCCTTTCTCGCA |
| SLC1A5 (ASCT2) | AGGCTTTCTCTGGCTGGTAAC | TCTTAGGTCCGGGAGGGTG |
| SLC2A1 (GLUT1) | CACTGTCGTGTCGCTGTTTG | AAAGATGGCCACGATGCTCA |
| SLC7A1 (CAT1) | CCGCCGGCTTGGATTCTGA | TTTGCACCCCATGTTGCTGT |
| SLC7A5 (LAT1) | CCGAGGAGGCAGCCAAG | TTGGACACATCACCCTTCCC |
| SLC16A1 (MCT1) | TGCGTGGGTACTGGAACAAG | TGCAGGTCAAATCCAAATATCGTT |
| SLC16A2 (MCT8) | CCTTCACCAGCTCCCTAAGC | ATGGCTGAAAGGCGAAGGAA |
| SLC22A1 (OCT1) | AATGCTGAGCTGTACCCCAC | CCCAACACCGCAAACAAAATG |
| SLC22A2 (OCT2) | AATCTCTACCCGCCTCCCTT | CACAGAGCTCGTGAACCAGT |
| SLC22A4 (OCTN1) | CAACGCCTTCAGCCTGTTTC | CTACGGGTGATGACAGCGTT |
| SLC22A5 (OCTN2) | TGTCCACCATTGTGACCGAG | GCAGGCTTCTTTCCCATCCT |
| SLC22A8 (OAT3) | TAGGACAGAGCAGGGACCTC | GAGAAGGTCATGGCACTGGG |
| SLC15A1 (PEPT1) | CTGTGGCGAAGTGGTCTTCT | GAATGTACTCGGCCCACTGT |
| SLC15A2 (PEPT2) | GCCTTCAGCAGCCTCTGTTA | ATGGCCAAGCACATACACCA |
| SLC38A2 (SNAT2) | CTCCTACCCCACCAAGCAAG | CCCACAATCGCATTGCTCAG |
| SLC38A3 (SNAT3) | AACCGCGAGGCCAGACATC | AGCAGCCCCTCTGAGTGTTT |
| SLC51A (OSTα) | TAAAGCTTGACCCCAGGTACA | ATGCTAGTGAGGGCAAGTTCC |
| SLC51B (OSTβ) | AGCATCCAGGCAAGCAGAAAA | GTGATCCTTGGCCTCATCCA |
| SLCO1A2 (OATP1A2) | AGCGTTCCAGGTATTTTTGTAATG | TCCCATGTTGCTCTTCAGGG |
| SLCO2B1 (OATP2B1) | CTGGAGCTCCCACCGTTATT | CTCACCCGCTGGCCCTATC |
| TfR | TCGTGTCCTCCCTTCATCCT | ACACAGAAGAACCTGCAGCC |
| β2-microglobulin (B2M) | GGCTATCCAGCGTACTCCAA | ACCAGTCCTTGCTGAAAGACAA |
| β-Actin | AACTACCTTCAACTCCATCA | GAGCAATGATCTTGATCTTCA |
